# Isolation of secreted proteins from *Drosophila* ovaries and embryos through *in vivo* BirA-mediated biotinylation

**DOI:** 10.1101/694091

**Authors:** Leslie M. Stevens, Yuan Zhang, Yuri Volnov, Geng Chen, David S. Stein

## Abstract

The extraordinarily strong non-covalent interaction between biotin and avidin (kD = 10^-14^-10^-16^) has permitted this interaction to be used in a wide variety of experimental contexts. The Biotin Acceptor Peptide (BAP), a 15 amino acid motif that can be biotinylated by the *E. coli* BirA protein, has been fused to proteins of interest, making them substrates for *in vivo* biotinylation. Here we report on the construction and characterization of a modified BirA bearing signals for secretion and endoplasmic reticulum (ER) retention, for use in experimental contexts requiring biotinylation of secreted proteins. When expressed in the *Drosophila* female germline or ovarian follicle cells under Gal4-mediated transcriptional control, the modified BirA protein could be detected and shown to be enzymatically active in ovaries and progeny embryos. Surprisingly, however, it was not efficiently retained in the ER, and instead appeared to be secreted. To determine whether this secreted protein, now designated secBirA, could biotinylate secreted proteins, we generated BAP-tagged versions of two secreted *Drosophila* proteins, Torsolike (Tsl) and Gastrulation Defective (GD), which are normally expressed maternally and participate in embryonic pattern formation. Both Tsl-BAP and GD-BAP were shown to exhibit normal patterning activity. Co-expression of Tsl-BAP together with secBirA in ovarian follicle cells resulted in its biotinylation, which permitted its isolation from both ovaries and progeny embryos using Avidin-coupled affinity matrix. In contrast, co-expression with secBirA in the female germline did not result in detectable biotinylation of GD-BAP, possibly because the C-terminal location of the BAP tag made it inaccessible to BirA *in vivo*. Our results indicate that secBirA directs biotinylation of proteins bound for secretion *in vivo*, providing access to powerful experimental approaches for secreted proteins of interest. However, efficient biotinylation of target proteins may vary depending upon the location of the BAP tag or other structural features of the protein.

## Introduction

Originating with the pioneering studies of Casadaban, Silhavy, Beckwith and co-workers^1-5^, experimental strategies involving the generation of proteins that have been attached genetically to exogenous protein or peptide tags have had an enormous impact upon progress in biological disciplines including biochemistry, cell and developmental biology, genetics, microbiology and molecular biology. A variety of protein tags that can be visualized [e.g. β-Galactosidase and fluorescent proteins such as Green Fluorescent Protein (GFP)]^1-9^ have enabled analyses of protein expression, abundance, subcellular localization and topology *in vivo*. Other protein tags (e.g. Glutathione-S-Transferase, Maltose-Binding Protein)^10-14^ have facilitated isolation of proteins-of-interest by affinity chromatography, thus permitting analyses of their structure, modification, and interaction with other factors.

The large size of the protein tags of the types described above can alter the behavior of the proteins-of-interest to which they have been fused. Accordingly, a variety of small peptide tags that interact either with characterized antibodies (several tags comprising characterized epitopes)^15-17^, Streptavidin/Streptactin (SBP-tag, Strep-tag)^18-20^, Calmodulin (Calmodulin-tag)^21^, Nickel or Cobalt chelate (His-tag)^22-26^ or anion exchange resin (polyglutamate tag)^27, 28^ have been utilized for the detection or isolation of proteins of interest to which they have been fused. Over the course of time, many additional protein tags have been developed with various useful properties^29^.

Biotin (also referred to as vitamin B7, vitamin H, and coenzyme R) is synthesized by plants, most bacteria, and fungi and acts as an enzyme cofactor, playing a critical role in some carboxylation, de-carboxylation and trans-carboxylation reactions^30^. *Escherichia coli* contains a single biotinylated protein, the biotin carboxyl carrier protein (BCCP) subunit of the acetyl-CoA carboxylase^31, 32^ which plays a critical role in fatty acid biosynthesis and degradation^33^. Biotinylation of BCCP is mediated by the *E. coli* BirA protein^34^. The minimal region of BCCP required for BirA-mediated biotinylation was defined as a 75 amino acid stretch of the protein^30^. Phage display allowed the identification of a 15 amino acid peptide (AviTag or BAP Tag) that is unrelated to the site of biotinylation in BCCP, but which has served as a convenient target for *in vivo* biotinylation by BirA of other proteins to which it has been attached^35^. As in *E. coli*, biotinylated proteins are similarly rare in other organisms; mammals, for example, contain only four biotinylated proteins^36^, a feature that would serve to limit interference from endogenous proteins in the detection and analysis of proteins heterologously biotinylated by BirA.

The strength of the avidin:streptavidin/biotin interaction^37, 38^ and the rarity of endogenous biotinylated proteins have combined to make *in vivo* biotinylation of proteins of interest by BirA an especially useful tool for their detection, analysis and isolation^39^. In addition, co-expression of BAP-tagged proteins with BirA has provided a method for purifying the resulting biotinylated fusion protein together with other proteins with which it forms complexes^39, 40^. In an approach that is similar to chromatin immunoprecipitation (ChIP), which has been used extensively to identify DNA sequences bound by specific transcription factors (TFs), BirA-mediated biotinylation has also provided a useful tool for the study of protein:chromatin interactions. In ChIP, antibodies targeting a TF of interest are used for immunoprecipitation of fragments of chromatin with which the TF interacts. However, for TFs for which useful antibodies do not exist, an alternative approach has been to BAP tag the TF, then use immobilized avidin to purify chromatin fragments that have been bound by that TF^43-45^. BirA’s ability to attach biotin, as well as a ketone isostere of biotin, has enabled various approaches for labeling BAP-tagged proteins *in vivo*^46, 47^. Another development that has increased the versatility of this approach is the isolation of promiscuous versions of BirA (BirA*) that do not require the presence of the BAP tag sequence and will instead biotinylate proteins based on their proximity to the protein carrying the BirA* enzymatic activity (proximity labeling). This has led to novel proteomic approaches in which BirA*-tagged fusion proteins are used to biotinylate interacting proteins or proteins that reside within the same subcellular compartment, which can then be visualized and/or isolated and identified^48-50^.

Here we add to the versatility of the BirA tool kit by demonstrating that a secreted version of BirA bearing an endoplasmic reticulum (ER)-retention signal is capable of performing *in vivo* biotinylation of a BAP-tagged secreted protein in *Drosophila* ovarian cells and embryos. However, these studies also indicate that care needs to be taken in constructing the fusion proteins to ensure that the BAP sequence will be accessible to co-expressed BirA when the protein is in its native conformation *in vivo*.

## Results

### A secreted version of the *E. coli* BirA protein is expressed and active in *Drosophila* ovarian cells and in the embryo

In an effort to develop a simple and efficient method for the isolation of secreted proteins relying on the high affinity interaction between avidin and biotin, we initially generated a secreted version of *E. coli* BirA that was designed to be retained in the ER. A PCR based approach was used to generate a BirA construct comprising the amino terminal 20 amino acids of the *Drosophila* secreted serine protease Easter^51, 52^ corresponding to its signal peptide, followed by the entirety of the BirA open reading frame, with the addition of the four amino acids lysine-glutamic acid-glutamic acid-leucine (KEEL) at the C-terminus of the fusion protein. The Easter signal peptide has been shown to direct the efficient secretion of heterologous proteins to which it has been fused^53, 54^, while the amino acid sequence KEEL is required for correct localization in some *Drosophila* ER proteins^55, 56^. A DNA fragment that encodes the resulting fusion protein, secBirA, was then introduced into *pUAST*^57^ and *pUASp*^58^, which are P-element based expression vectors that can be expressed under the control of the yeast transcriptional activator Gal4 in somatic and germline-derived tissues of *Drosophila*, respectively.

Expression of genes cloned into *pUASp* under the control of the *Nanos-Gal4::VP16* driver element^58^ leads to the production of protein in the germline-derived ovarian cells (15 nurse cells and oocyte) and in the progeny embryo, respectively, while expression of genes cloned into *pUAST* under the control of the *CY2-Gal4* driver element leads to the production of protein in the ovarian follicle cells^59^. Ovarian and embryonic protein extracts from females carrying *pUASp-secBirA* together with *Nanos-Gal4::VP16*, and extracts from ovaries of females bearing *pUAST-secBirA* together with *CY2-Gal4*, were subjected to Western Blot analysis using an antibody directed against BirA (Creative Diagnostics). Two specific bands were detected (Figure 1) that are likely to correspond to full-length secBirA bearing the signal peptide and secBirA from which the signal peptide has been cleaved, although these bands are somewhat smaller in apparent molecular weight than expected for these proteins, 38 kD and 36 kD, respectively. These results indicate that secBirA protein was successfully expressed in ovaries and embryos under Gal4-mediated transcriptional control. secBirA expression in female flies did not lead to any detectable perturbations of oogenesis or embryogenesis.

**Fig 1.**
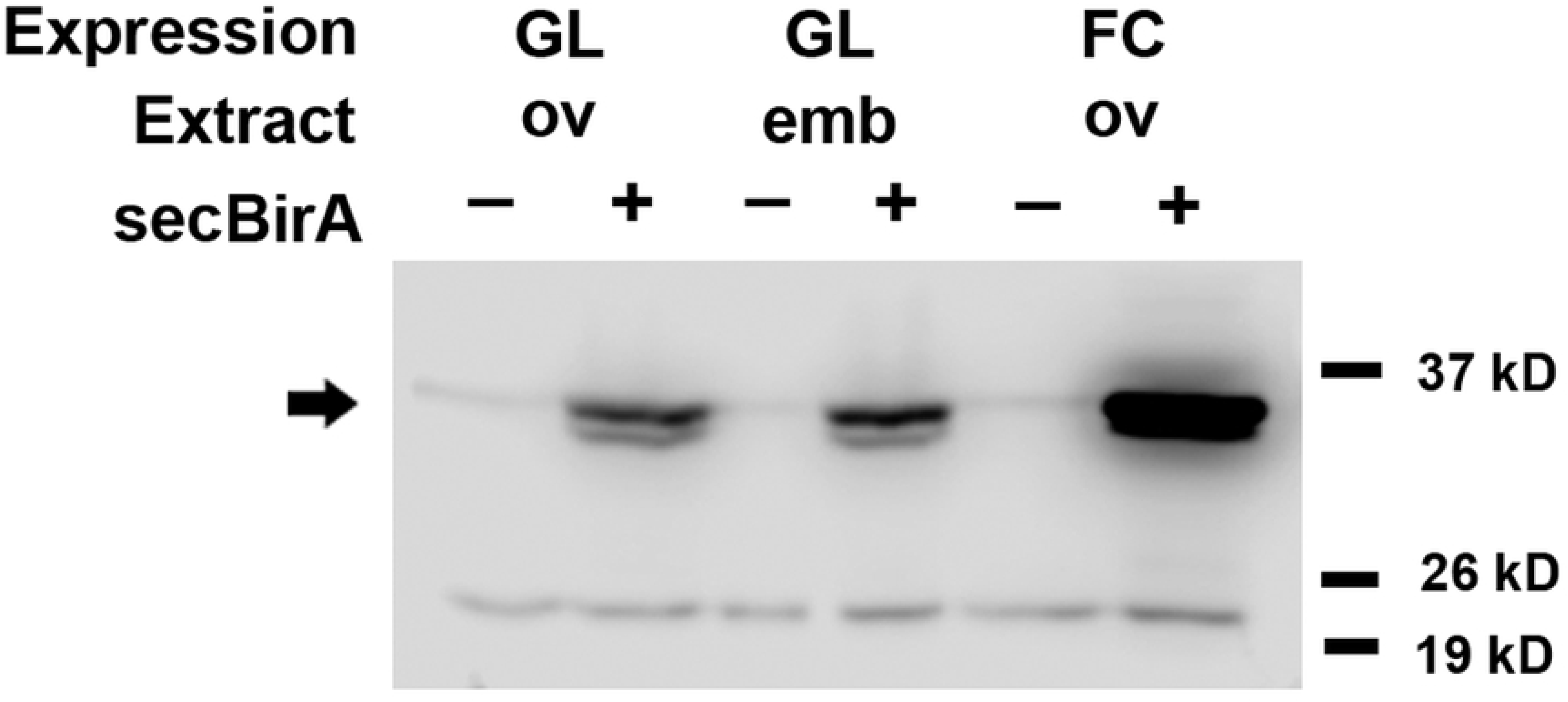
Gal4-mediated expression of secBirA protein in *Drosophila* ovaries and embryos. Protein homogenates were prepared from ovaries (GL ov) and progeny embryos (GL emb) from females expressing secBirA(+) in the female germline (nurse cells) under the control of the *Nanos-Gal4::VP16* driver element^58^ and from homogenates isolated from the ovaries of females expressing secBirA(+) in the follicle cell layer (FC ov) under the control of the *CY2-Gal4* driver element^59^. These were subjected to Western blot analysis using an antibody directed against full-length BirA protein. Control homogenates were from ovaries and embryos carrying the corresponding Gal4 driver elements in the absence of the secBirA expression construct (-). Note the presence of a pair of specific bands (indicated by the arrow) in homogenates from tissues expressing secBirA, whose molecular masses correspond closely with the predicted molecular weights of full-length secBirA and of secBirA from which the signal peptide has been cleaved.

To examine the subcellular localization of secBirA, we carried out whole mount immunohistochemical staining and confocal imaging of ovaries from *CY2-Gal4*/*pUAST-secBirA* females and of embryos from *pUASp-secBirA*/*nanos-Gal4::VP16* females, both of which also expressed a GFP-tagged version of Protein Disulfide Isomerase (PDI-GFP)^60^. Consistent with its ER localization, PDI-GFP-associated fluorescence exhibited a reticular distribution in both ovarian follicle cells (Figure 2A) and in the syncytial blastoderm embryo (Figure 2D). Surprisingly, secBirA did not co-localize extensively with PDI-GFP in either ovarian follicle cells (Figure 2B, C), or in the embryo (Figure 2E, F). However, secBirA did appear to undergo secretion into the cleft between the apical surface of the follicle cells and the developing oocyte (see arrow, Figure 2B). In embryos, secBirA exhibited a punctate distribution in the cytoplasm (Figure 2E, F). It was not possible to determine whether secBirA was secreted into the perivitelline space lying between the embryo plasma membrane and the inner vitelline membrane (VM) layer of the eggshell, as the whole mount staining protocol requires removal of the VM, which leads to the loss of proteins that have been secreted into the perivitelline space.

**Fig 2.**
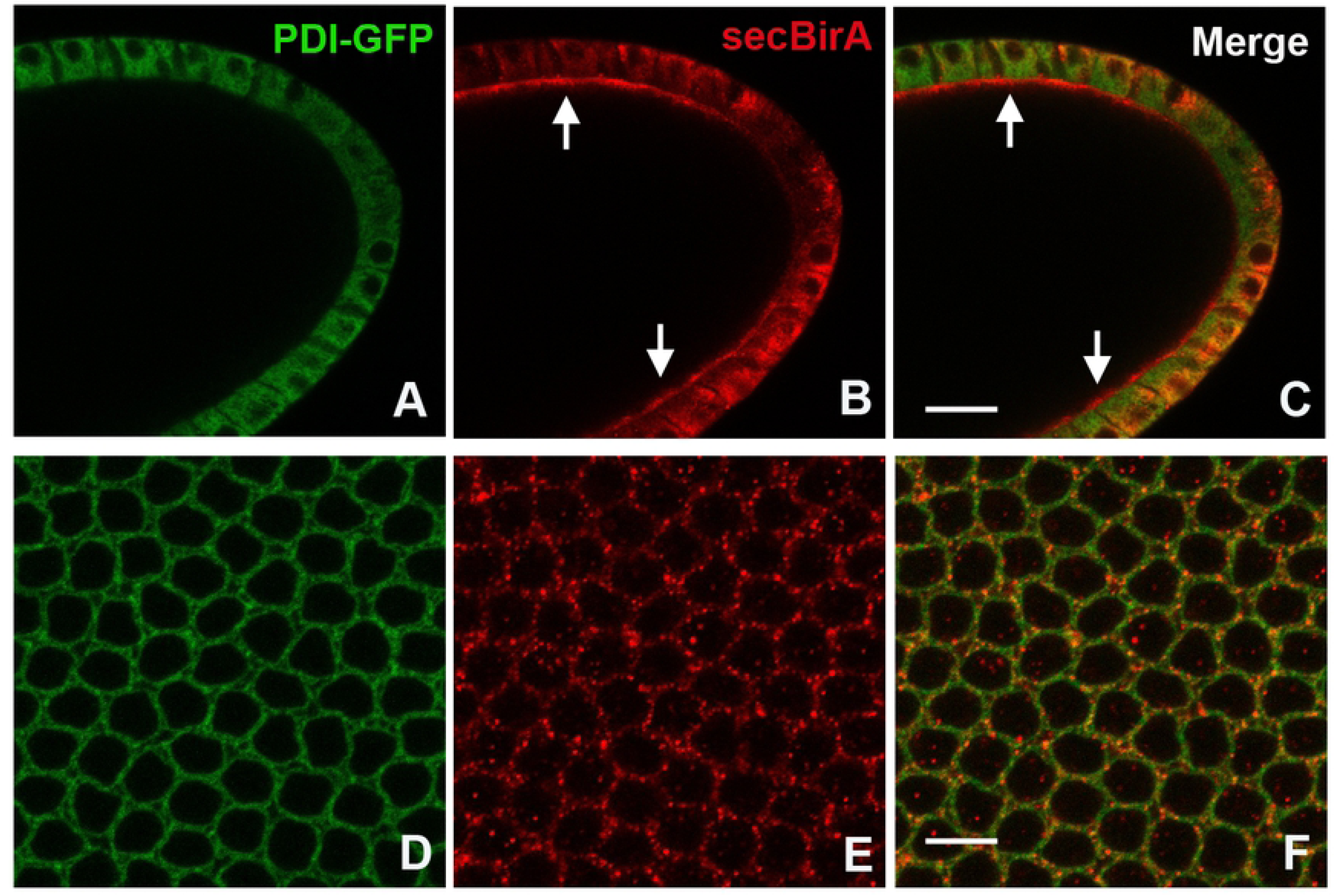
Visualization of the distribution of secBirA expressed in *Drosophila* ovaries and embryos. Confocal optical sections of the follicle cell layer surrounding the oocyte of a stage 10 egg chamber (A, B, C) and of the surface view of a syncytial blastoderm stage embryo (D, E, F) expressing the ER marker PDI-GFP^60^ (A, D) and stained with anti-BirA (B, E). (C) and (F) show merged images of the PDI-GFP expression and secBirA staining patterns in the follicle cell layer and the embryo, respectively. Note that PDI-GFP, which is known to be localized to the ER, and secBirA do not exhibit extensive colocalization. Although secBirA bears an ER-retention signal it appears to be secreted, as shown by its presence in the perivitelline space between the apical surface of the follicle cells and the large developing oocyte [see arrows in (B), (C)]. secBirA exhibits a punctate distribution in the syncytial blastoderm embryo. The scale bars represent 22.5 microns in (C) and 7.5 microns in (F).

To confirm that the secBirA expressed in ovarian cells and in the embryo was functional, we assayed for biotin ligase activity associated with its expression. Protein extracts were prepared from ovaries and embryos derived from females that expressed BirA in the germline and from the ovaries of females that expressed it in the follicle cells. As controls, extracts were also prepared from tissues derived from females carrying only the Gal4 drivers. Homogenates were prepared in activity assay buffer and given as a substrate Maltose Binding Protein carrying the BAP Tag (MBP-BAP). After the reaction was complete it was loaded onto an SDS-PAGE gel and blotted and processed for biotin detection. As shown in Figure 3, ovarian extracts expressing secBirA in either the germline or the follicle cells, as well as embryonic extracts containing secBirA, were able to transfer biotin to MBP-BAP. In the absence of expressed secBirA, no MBP-BAP biotinylation was detected.

**Fig 3.**
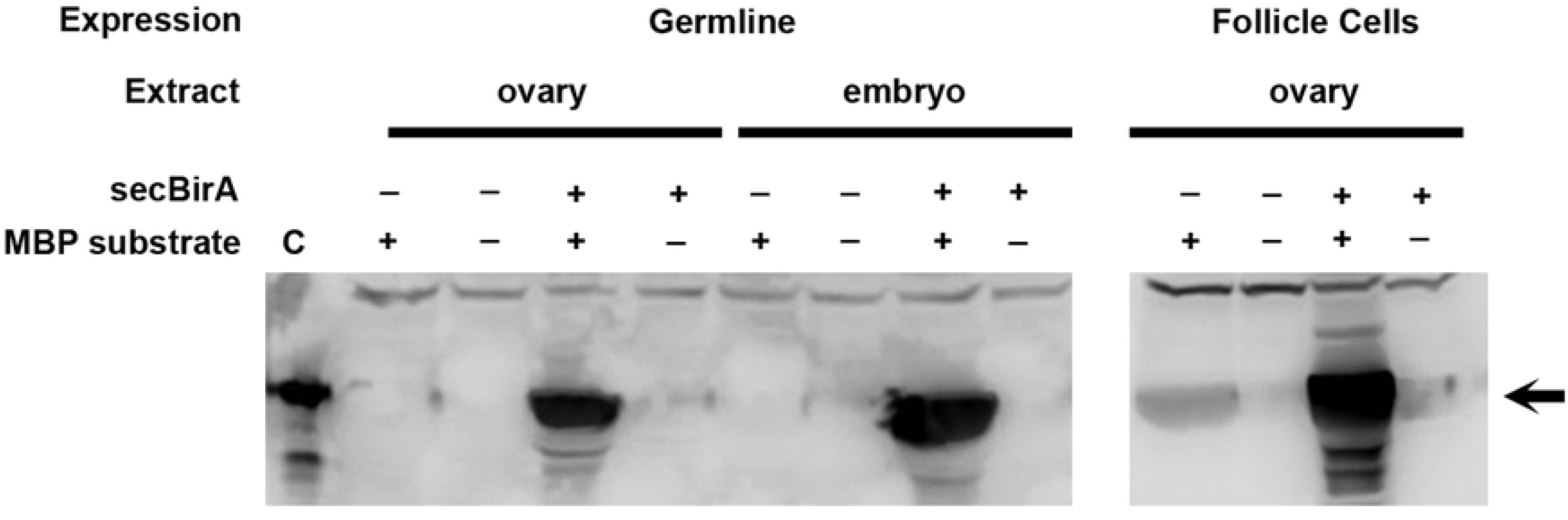
secBirA expressed in *Drosophila* ovaries and embryos exhibits biotin ligase activity. Protein extracts from ovaries and embryos expressing (+) or lacking (-) secBirA were incubated with (+) or in the absence of (-) a BirA substrate, Maltose Binding Protein fused in-frame to the Biotin Acceptor Peptide (MBP substrate). After SDS-PAGE and blotting to nitrocellulose, biotinylated MBP was detected with streptavidin-HRP. The lane at far left (labelled C) contains commercially available (Avidity) biotinylated MBP (bio-MBP) as a positive control. When incubated together with substrate, extracts from ovaries and progeny embryos of females expressing secBirA in the germline, and extracts of ovaries from females expressing secBirA in the follicle cell layer, exhibited a strong signal corresponding to biotinylated MBP (see arrow).

### BAP-tagged versions of the *Drosophila* patterning proteins Torsolike and GD are functional

Despite the apparent inability of the C-terminal KEEL motif to mediate efficient retention of secBirA in the ER, the observation that secBirA undergoes secretion suggested that the protein would nevertheless co-reside with potential target proteins during their transit through the secretory pathway and might therefore be able to catalyze their biotinylation. We selected *Drosophila* Torsolike (Tsl)^61-63^ and Gastrulation Defective (GD)^64-66^ as secreted proteins that could potentially serve as substrates for secBirA-mediated biotinylation.

The Tsl protein participates in patterning along the anterior/posterior (AP) axis of the developing embryo. Specifically, Tsl is required for the formation of the two termini, the acron at the anterior and the telson at the posterior^61-63^. The *tsl* gene is expressed in two subpopulations of follicle cells adjacent to the anterior and posterior ends of the developing oocyte^62, 63^. The protein product is secreted from those cells and becomes localized to the polar regions of the VM layer of the eggshell^67^ as well as to the plasma membrane at the two ends of the embryo^62, 68^. The polar localization of Tsl is required to mediate spatially-restricted activation of the receptor tyrosine kinase Torso at the two termini of the developing embryo, which is necessary for proper AP patterning^69-70^. Expression of *tsl* in all follicle cells results in embryos that are terminalized^62, 63, 72^. The formation of segments is suppressed while terminal elements expand, typically leading to an embryo bearing few cuticular pattern elements aside from two large fields of Filzkörper (tracheal spiracle) material.

The GD protein participates in patterning of the *Drosophila* embryo along the dorsal-ventral axis^64, 65^. The *gd* gene is normally transcribed in the nurse cells of the ovary^66^ and the protein product is present in the perivitelline space of the egg^73, 74^, where it participates in a proteolytic cascade^75-77^ that results ultimately in the formation of the active ligand for the Toll receptor ventrally within the perivitelline space^78-80^. Ventral activation of plasma membrane-localized Toll receptor^81^ by its ligand is necessary for the correct formation of the embryonic DV axis. Transgene-mediated overexpression of GD in the germline leads to the formation of ventralized embryos with an expansion of ventral pattern elements^73^. Typically, cuticles formed by these ventralized embryos exhibit ventral denticles all around their DV circumferences, and lack dorso-laterally derived Filzkörper material altogether.

We used high fidelity PCR to generate DNA clones encoding Tsl and GD that carried at their C-termini 2 glycine residues followed by the 15 amino acid long BAP tag^35^, which we refer to as Tsl-BAP and GD-BAP, respectively. The DNA clone encoding Tsl-BAP was introduced into pUAST^57^ while *GD-BAP* was introduced into pUASp^58^. As has been previously observed for wild-type tsl^62, 63, 72^, expression of *tsl-BAP* throughout the follicle cell layer (Figure 4B) led to the formation of embryos comprised solely of expanded terminal pattern elements. Similarly, as has been seen previously for wild-type *gd*^73^, transgenic overexpression of *gd-BAP* in the female germline led to the formation of progeny embryos that were ventralized (Figure 4C). Accordingly, we conclude that Tsl-BAP and GD-BAP retained function.

**Fig 4.**
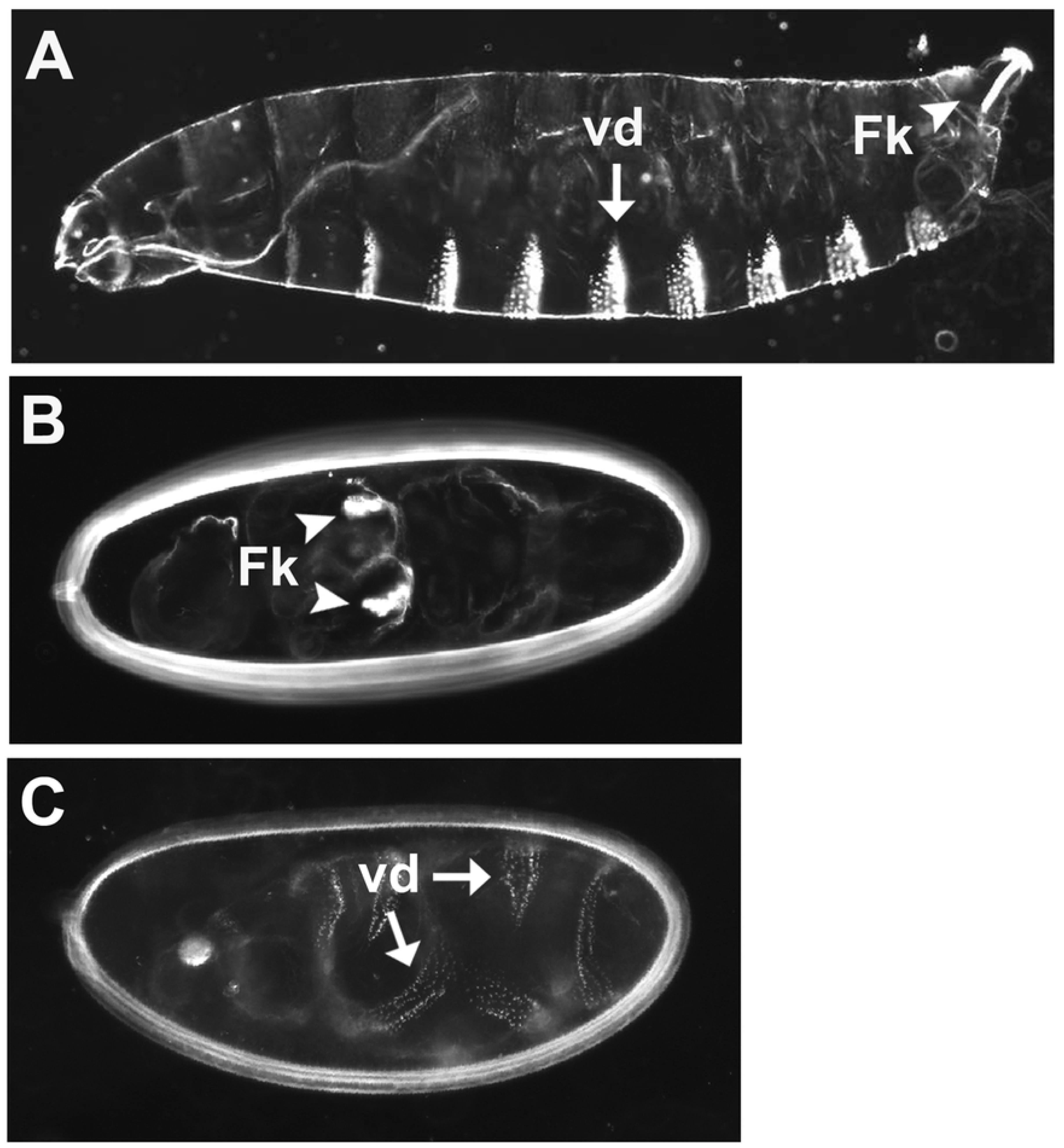
BAP-tagged versions of the Tsl and GD proteins are functional. (A) Wild-type first instar larval cuticle showing ventral denticle bands (vd, arrow) and Filzkörper (Fk, arrowhead) structures. (B) Embryonic cuticle produced by a mother expressing Tsl-BAP throughout the follicle cell layer under the control of the *CY2-Gal4* driver element. As seen previously with wild-type Tsl^62, 63, 72^, uniform expression in the follicle cell layer leads to the formation of progeny embryos that are made entirely of terminal pattern elements. The segmented regions of the embryo are suppressed, with the loss of ventral denticles and the presence of patches of Filzkörper material (Fk, arrowheads) that have formed in the central region of the embryo. (C) Embryonic cuticle produced by a mother expressing *GD-BAP* mRNA in ovarian nurse cells under the control of the *Nanos-Gal4::VP16* driver element. As has been seen for wild-type GD protein^73^, transgenic overexpression in the female germline leads to the formation of ventralized embryos. Pattern elements that are normally found in dorsal and dorsolateral regions of the larvae, such as Filzkörper, are absent, while there is an expansion of ventral structures such as ventral denticles (vd, arrows) around the DV circumference of the embryo.

### secBirA can biotinylate secreted proteins in *Drosophila*

To test whether GD-BAP or Tsl-BAP can be biotinylated by secBirA *in vivo*, we co-expressed the BAP-tagged proteins with secBirA in either the germline (GL) under the control of *nanos-Gal4::VP16*^58^ or in the ovarian follicle cell layer (FC) under the control of *CY2-Gal4*^59^. Protein extracts from either ovaries or progeny embryos were then subjected to SDS-PAGE and Western blotting followed by biotin detection (Fig. 5). A conspicuous band with an apparent molecular weight corresponding to that of Tsl-BAP was observed in extracts of both ovaries (lanes 5 and 6) and embryos (lane 7) from females expressing Tsl-BAP and secBirA in their follicle cells. However, no band of a molecular weight consistent with that of biotinylated GD-BAP, either its ∼42 kD processed form, or the unprocessed form of 57kD or 61 kD (depending upon signal peptide removal), was detectable in extracts of ovaries in which GD-BAP was co-expressed with secBirA in the germline (lane 2). This was surprising, as endogenous GD protein is expressed in the germline and, as noted above, GD-BAP expressed under the control of *nanos-Gal4::VP16* is active in DV patterning. To test whether the failure to detect biotinylation of GD-BAP was related to the germline milieu, we expressed it together with secBirA in the follicle cells, where we had shown BirA to be active on Tsl-BAP. However, GD-BAP was not detectably biotinylated in the follicle cell layer, either (lane 3). In contrast to GD-BAP, a different version of BAP-tagged Tsl that was cloned into *pUASp* was biotinylated when co-expressed in the germline with secBirA (data not shown). As secBirA exhibited activity on MBP-BAP in both germline and follicle cell-derived extracts, and Tsl-BAP was biotinylated when expressed in both types of tissues, the failure to detect biotinylation of GD-BAP seems likely to be related to the location of the BAP tag within that particular fusion protein. Finally, it should be noted that we consistently detected a set of high molecular weight proteins that exhibit endogenous biotinylation that is not dependent upon the BAP transgenes (Fig. 5) or the expression of secBirA (data not shown).

**Fig 5.**
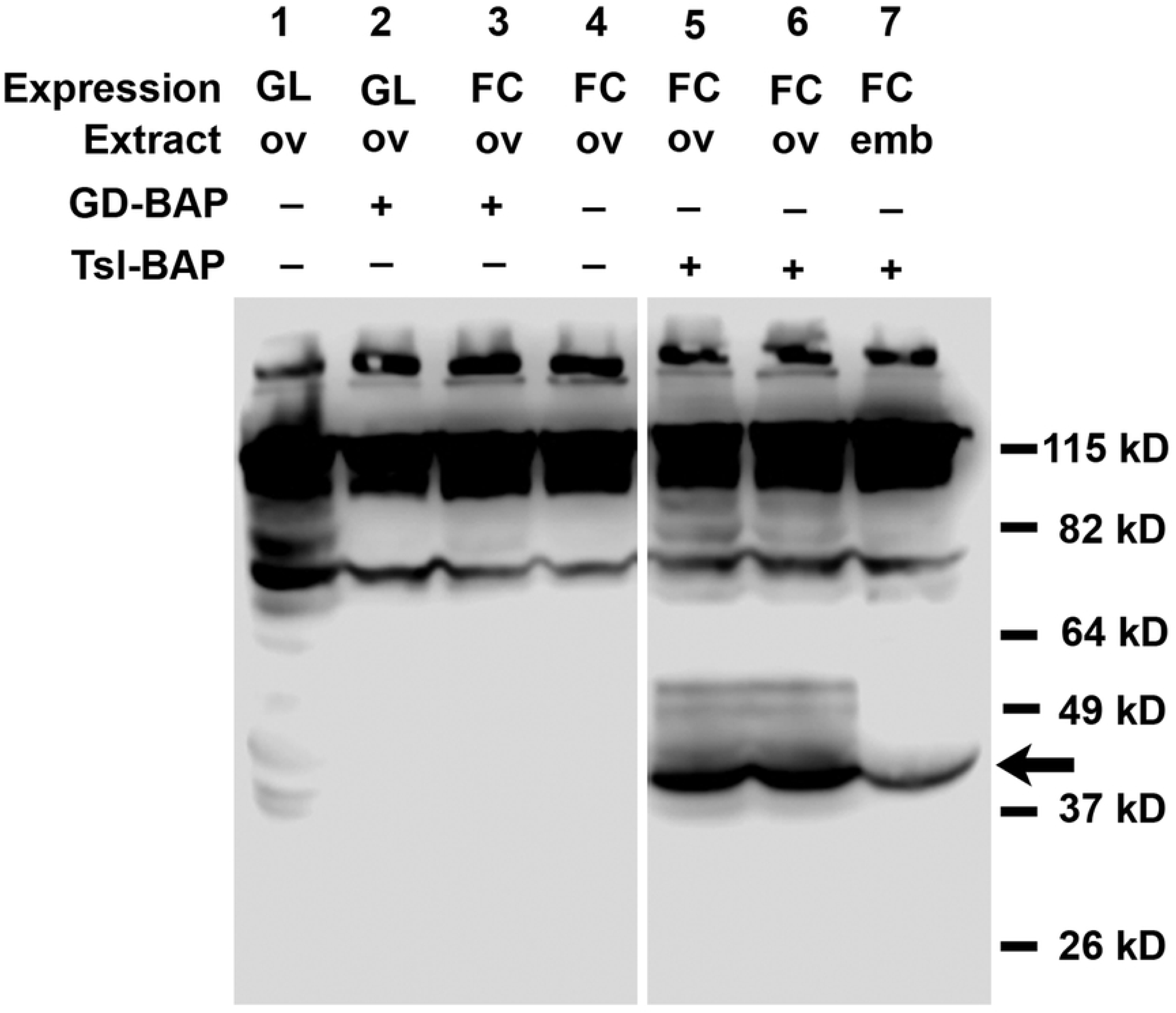
BAP-tagged Tsl is detectably biotinylated when co-expressed with BirA but GD is not. Protein extracts from ovaries (ov) or embryos (emb) derived from females expressing secBirA in the germline (GL) or follicle cell layer (FC) either alone (-) or co-expressed (+) with Tsl-BAP or GD-BAP. Biotinylated Tsl-BAP is indicated by the arrow. No biotinylated bands near the expected sizes of either unprocessed or processed GD-BAP, 58.7 kD and approximately 42 kD, respectively, are detectable. Both panels are derived from the same gel and blot; two irrelevant lanes in between were removed from the figure for clarity. 150 μg of protein was loaded into each lane.

### secBirA permits enrichment of biotinylated secreted proteins in *Drosophila*

The experiments outlined above indicate that secBirA can perform *in vivo* biotinylation of Tsl-BAP. The ability to perform affinity purification or enrichment of secreted biotinylated BAP-tagged proteins would significantly extend the usefulness of this approach. To explore this possibility, we utilized Streptavidin coupled to magnetic beads (Thermo Fischer Scientific) to carry out a small-scale batch affinity isolation of Tsl-BAP from extracts of embryos produced by females co-expressing secBirA and Tsl-BAP in the follicle cell layer under the control of CY2-Gal4. As a negative control, extracts of embryos from females expressing secBirA in the absence of Tsl-BAP were subjected to the same isolation protocol. Aliquots were taken at various stages of the procedure, subjected to SDS-PAGE and Western blotting, and then processed for biotin detection. Figure 6 demonstrates that biotinylated Tsl-BAP was specifically detected in the starting material (SM) homogenate and the post-equilibration (PE) sample that was obtained following column chromatography through G-25 Sepharose to remove free biotin. The post-binding (PB) supernatant that was removed following overnight incubation with the streptavidin resin was largely devoid of biotinylated Tsl-BAP, indicating that binding of the biotinylated protein to the resin was highly efficient. After extensive washing the bound protein was eluted from the resin and the resulting eluate (E) exhibited a strong signal of biotinylated Tsl-BAP (Fig. 6). The absence of a biotinylated protein of similar molecular weight to Tsl-BAP in the negative control confirms the identification of this band as corresponding to Tsl-BAP. This finding demonstrates the utility of *in vivo* biotinylation by secBirA as a method for tagging secreted proteins with an element that permits their enrichment from complex mixtures.

**Fig 6.**
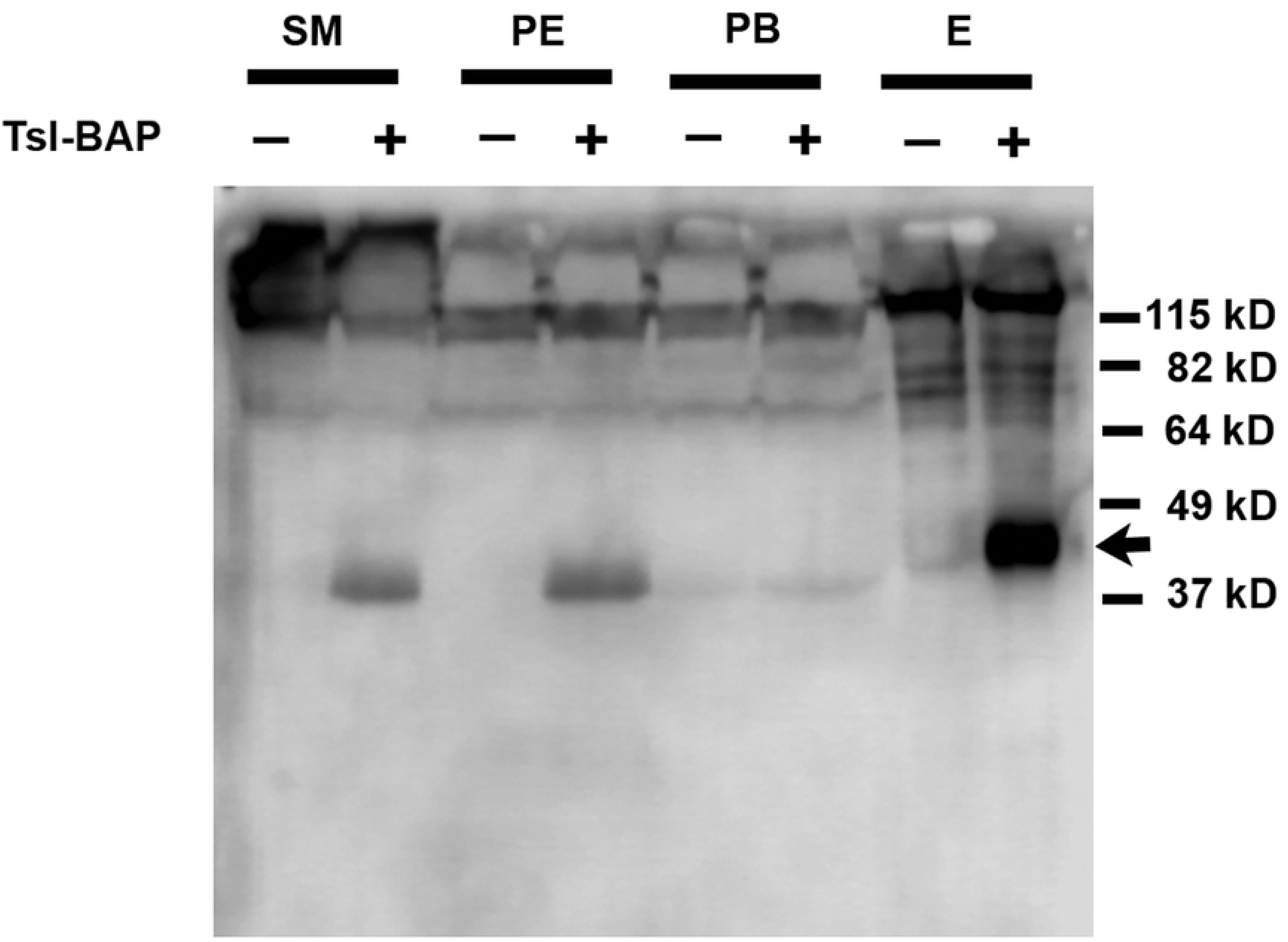
Tsl-BAP can be isolated from extracts using streptavidin-coupled resin. A procedure to isolate biotinylated proteins was carried out using extracts from embryos derived from females expressing secBirA and Tsl-BAP in the follicle cell layer (+), or from sibling females carrying only the secBirA transgene (-). The samples shown are the starting material (SM), the post-equilibration sample (PE), the post-binding supernatant (PB) after incubation with streptavidin resin, and the sample eluted (E) from the resin after extensive washing steps. The samples were subjected to SDS-PAGE, blotted and processed for biotin detection. 100 μg of protein were loaded onto the SM, PE and PB lanes. For the eluate lanes, 2 μl of a total volume of approximately 50 μl was loaded. Biotinylated Tsl-BAP is detected in the SM and PE samples, but is absent from the PB samples, indicating that biotinylated Tsl-BAP was effectively removed from the mixture by the streptavidin resin.

The eluate lane in Figure 6 represents 4% of the total eluate that was obtained from an initial quantity of embryos corresponding to 350 mg. 80% of the eluates from the experimental and negative control isolations were loaded onto a separate gel that was stained using Coomassie Brilliant Blue to visualize protein bands. Although many protein bands were detected, no band in the vicinity of the expected Tsl-BAP size range appeared to be specific to the experimental sample (data not shown). Moreover, despite extensive washing during the procedure, numerous protein bands corresponding to non-biotinylated contaminants were present in both the control and experimental samples. Thus, although this approach may be useful for several experimental tests (of protein processing, modification, and interaction with candidate proteins for which antibodies are available), additional measures to increase the specificity of the isolation would be necessary to address other experimental questions (e.g. mass spectrometry of interacting proteins, structural studies), as discussed below.

## Discussion

*In vivo* biotinylation of proteins by *E. coli* BirA has proven a useful technology in a number of experimental applications^39, 43-50^. The results described above indicate that for some secreted proteins expressed in *Drosophila* tissues, co-expression of a BAP-tagged variant together with secBirA can provide an effective means of biotinylation for detection and isolation. *Drosophila* was considered to be an ideal model organism for this approach because a major component of most *Drosophila* food recipes is yeast, an excellent source of biotin. Moreover, *Drosophila* food can easily be supplemented with additional biotin. Although our studies have focused on proteins expressed in the ovary and embryo, the use of this approach is not restricted to those tissues. Numerous *Drosophila* stocks expressing Gal4 in a wide variety of tissue specific patterns are available. These Gal4 driver lines can be used to direct the expression of pUAST-secBirA, together with transgenic UAS-directed BAP-tagged versions of other secreted proteins-of-interest for the analysis of biotinylated protein. Similarly, co-transfection of a Gal4 expression plasmid, pUAST-secBirA, and a UAS-driven transgene encoding a BAP-tagged secreted protein-of-interest into cultured *Drosophila* cells could also facilitate the isolation of the protein-of-interest for further analysis.

For *in vivo* biotinylation by BirA to succeed, it is essential that BirA and the substrate co-reside, at least transiently, in the same cellular compartment. Both Tsl and GD are known to be secreted proteins bearing N-terminal signal peptides. Although, as discussed below, the secBirA in this study was designed to be retained in the endoplasmic reticulum, immunohistochemical staining of ovaries in which secBirA was expressed in the follicle cell layer suggested that secBirA was being secreted by the follicle cells and thus was likely to be transiting through the secretory pathway along with its target proteins. The biotinylation of Tsl-BAP co-expressed with secBirA in either the germline or the follicle cells is consistent with this hypothesis. It was surprising, therefore, that biotinylation of GD-BAP was not detected following its co-expression with secBirA in either the germline or the follicle cell layer. As the GD-BAP transgene did produce protein that was capable of providing GD function in embryonic patterning, the most likely explanation for this discrepancy is that the carboxy terminus of GD, and/or the BAP tag added at that location, is not exposed at the protein surface under native conditions, making it inaccessible as an enzymatic substrate for BirA. The determination of the three-dimensional structure of GD could confirm or refute this possibility. The likelihood that protein conformation influences the efficiency of *in vivo* biotinylation of a target protein by BirA highlights the need for care in selecting the location at which the BAP tag is inserted into the protein-of-interest.

As GD is known to be present in complexes with other secreted members of the “dorsal group” of maternal effect proteins controlling DV polarity^74^, an alternative explanation is that those protein/protein interactions interfere with the biotinylation of GD-BAP by BirA. The ventralized phenotype produced by expression of GD-BAP, however, suggests that the protein is present at very high levels that would likely result in some uncomplexed GD-BAP. Nevertheless, this potential complication is another issue that should be considered with respect to the placement of the BAP tag or even whether *in vivo* biotinylation is a viable option for a particular protein.

The four amino acids KDEL, or some close variant of this sequence, present at the carboxy terminus of proteins translated into the secretory compartment, is the canonical target of the KDEL receptor, which is responsible for retrieval of ER that have trafficked to the Golgi Apparatus^82, 83^. In *Drosophila*, the presence of the amino acid sequence KEEL at the carboxy terminus is also a signal for retrieval of proteins from the Golgi^56^. Among ER-localized proteins bearing C-terminal KEEL sequences are Windbeutel^84^ and Seele^85^, which participate in embryonic patterning and whose functions require that they be expressed in the ovarian follicle cells^55, 78^ and in the female germline^85, 86^, respectively. For this reason, we elected to include the KEEL sequence at the C-terminus of secBirA as a means of directing ER retention. We were therefore surprised to observe that while secBirA is secreted, it does not exhibit extensive colocalization with PDI-GFP, a known resident of, and useful marker for, the endoplasmic reticulum^60^. Thus, secBirA does not appear to be efficiently retrieved back to the ER. We suspect that the presence the KEEL sequence present at the C-terminus of the protein does not lead to efficient binding by the KDEL receptor protein.

It is unclear whether an exposed C-terminal KEEL motif is sufficient to direct ER localization in all KDEL receptor-retrieved proteins in *Drosophila*, or whether other determinants are required for ER retention. If the failure of secBirA to be retained in the ER results solely from a lack of availability of the KEEL sequence to interact with the KDEL receptor, then adding KEEL to BirA in a context that renders the KEEL exposed might facilitate retention of the protein in the ER and thereby increase the efficiency of biotinylation of BAP-tagged secreted proteins. In unpublished work we have generated a transgenic version of the Seele protein fused to GFP, in which the KEEL sequence was moved from the carboxy end of Seele to that of GFP. Like native Seele, this Seele-GFP localizes to the ER, and can substitute for the endogenous Seele protein in DV patterning. Therefore, as a C-terminal fusion to GFP, the KEEL motif appears to be capable of engaging with the KDEL receptor. Therefore, if a surface exposed KEEL motif is the sole determinant necessary for ER retention of secreted proteins, then attachment of GFP-KEEL to the C-terminus of secBirA (i.e. secBirA-GFP KEEL) may facilitate efficient ER localization and improve the efficiency of biotinylation of secreted proteins by this protein, providing the attachment of GFP to the C-terminus of BirA does not perturb its enzymatic activity.

As demonstrated in Figures 5 and 6 and data not shown, several endogenous high molecular weight biotinylated proteins reside in *Drosophila* ovaries and early embryos. For applications in which the target can be subjected to SDS-PAGE and excised from a gel, or otherwise separated from these endogenous biotinylated proteins, their presence may not be problematic. For other applications, however, they are likely to generate a large signal that may overwhelm that of the target. One example would be histochemical visualization of the abundance and subcellular distributions of BAP-tagged biotinylated proteins in living tissues. Similarly, if the ultimate goal were to identify proteins that interact with the target by using the streptavidin-biotin reaction to pull the complexes out of a cellular extract and then subject them to mass spectrometry, the presence of so many contaminating proteins and complexes would likely be prohibitive.

One approach to mitigate this problem is to add an additional tag, such as hexa—histidine(his6), that permits an orthogonal purification method to be applied to the fusion protein. The tandem application of affinity purification protocols for polyhistidine and biotinylation tags has been successfully applied to isolate ubiquitinated proteins for mass spectrometric analysis^87^. In our experimental system this approach would be expected to result in a relatively pure preparation of the target protein without contaminating endogenous biotinylated proteins. The his6 tag has another important feature, which is that metal chelate affinity chromatography of his6-tagged proteins, like the biotin/strept(avidin) interaction, can be performed under strongly denaturing as well as non-denaturing conditions. This would allow the tandem purification protocol to be used in experiments to identify proteins that interact with the target protein, but for which strong denaturing conditions must be used, following a protein crosslinking step, for example, to ensure that the interacting proteins do not dissociate from one during the affinity purification. In preliminary experiments, we have shown that biotinylated Tsl-BAP can be isolated from extracts generated from embryos exposed to the crosslinking agent paraformaldehyde at a concentration of 1% for 12 or 20 minutes, comparable to^88-91^, suggesting that the BAP tag may be useful for this purpose.

In summary, the generation of secBirA and the demonstration that it is functional and can be used to biotinylate selected secreted proteins in *Drosophila* adds to the repertoire of experimental approaches that can be used to examine protein structure and function in *Drosophila*. With additional optimization (e.g. the addition of a second affinity tag) the experimental versatility of this approach should expand yet further.

## Materials and Methods

### *Drosophila* Stocks and Maintenance

All stocks were maintained employing standard conditions and procedures. Transgenic lines were generated in a *w*^1118^/*w*^1118^ mutant derived from the OregonR strain. The strain carrying the *nanos-Gal4::VP16* insertion was a kind gift of Dr. Pernille Rørth^58^. The strain carrying the *CY2-Gal4* insertion^59^ was a kind gift of Dr. Trudi Schüpbach.

The strain expressing PDI-GFP is described in Bobinec et al., 2003^60^.

### Examination of Embryonic Phenotypes

For the examination of embryonic phenotypes, larval cuticles were prepared according to Van der Meer (1977)^92^.

### DNA constructs

secBirA, a secreted derivative of the *E. coli* biotin ligase BirA, carries at its amino terminus the N-terminal 22 amino acids of the Easter protease, including the Easter secretory signal peptide, followed by the full-length BirA protein, with the 4-amino acid sequence KEEL, which has been shown to act as an ER retention signal in *Drosophila*^56^, located at the carboxy terminus of the protein. For the construction of *Drosophila* expression vectors encoding secBirA, the two DNA oligonucleotides: 5’-CACCAAAATGCTAAAGCCATCGATTATCTGCCTCTTTTTGGGCATTTTGGCGAAATCATCGGCGGGCCAGTTCATGAAGGATAACACCGTGCCACTG-3’ and 5’-CTATTATCACAGTTCCTCTTTTTCTGCACTACGCAGGGATATTTCACCGCCCATCC AGGG-3’ were employed in a high-fidelity PCR reaction (Q5® High Fidelity DNA Polymerase, New England Biolabs) using a plasmid bearing the *E. coli BirA* gene as template. The resulting DNA fragment was gel purified and introduced into the Gateway® Entry Vector, pENTR™ by Topoisomerase I based directional ligation (Invitrogen™ cat. #K240020), yielding plasmid *pENTR-secBirA*. In vitro recombination using the Gateway® LR Clonase® II enzyme mix (Invitrogen cat. #11791-020) was then used to introduce the secBirA-encoding DNA sequences into the Gateway® Destination vectors, *pPW* and *pTW* (*Drosophila* Genomics Resource Center), yielding the expression clones *pUASp-secBirA* and *pUAST-secBirA*. *pPW* is a derivative of the *Drosophila* female germline/nurse cell-specific Gal4-dependent expression vector *pUASp*^58^, while *pTW* is a derivative of *pUAST*^57^, a Gal4-dependent expression vector that permits expression in somatic cells in *Drosophila*, including the ovarian follicle cells.

Tsl-BAP is a derivative of the terminal class protein Torso-like (Tsl) bearing the full-length Tsl open reading frame followed by a pair of glycine residues and finally the 15 amino acid long Biotin Acceptor Peptide (GLNDIFEAQKIEWHE) that is a substrate of BirA^35^, at its carboxy terminus. For the construction of a *Drosophila* expression vector encoding Tsl-BAP, the two DNA oligonucleotides: 5’-CACCAAAATGCGGTCGTGGCCTGGCC-3’ and 5’-TTATCACTCGTGCCACTCGATCTTCTGGGCTTCAAATATGTCATTCAAACCGCCTCCTCGGGTGGGATGACTCTGCGGCATGTTAAGC-3’ were utilized in a high-fidelity PCR reaction using a plasmid bearing a *Drosophila* cDNA encoding the *tsl* gene as template. The resulting DNA fragment was gel purified and introduced into the Gateway® Entry Vector, pENTR™ by Topoisomerase I based directional ligation, then recombined into the Gateway® Destination vectors, *pTW* as described above, yielding the expression clone *pUAST-Tsl-BAP*.

GD-BAP is a derivative of the dorsal group serine protease Gastrulation Defective (GD) bearing the full-length GD open reading frame followed by a pair of glycine residues and finally the 15 amino acid long Biotin Acceptor Peptide (GLNDIFEAQKIEWHE)^35^, which is the substrate of BirA, at its carboxy terminus. For the construction of a *Drosophila* expression vector encoding GD-BAP, the two DNA oligonucleotides: 5’-ACGTACGCGGCCGCAAAATGAGGCTGCACCTGGCGGCGATCC-3’ and 5’-ACGTACTCTAGACTACTCGTGCCACTCGATCTTCTGGGCTTCAAATATGTCATTCAAACCGCCTCCAATTACAAAGGCCGTGATCCAGTCCAGAAACTTGGCC were employed in a high-fidelity PCR reaction (Q5® High Fidelity DNA Polymerase, New England Biolabs) using a plasmid bearing a *Drosophila gd* cDNA as template. The resulting DNA fragment was gel purified, subjected to digestion with the restriction endonucleases Not I and Xba I, and subcloned into similarly digested *pUASp*^58^ to generate *pUASp-GD-BAP*.

Transgenic lines bearing the constructs described above were generated by conventional P-element mediate transformation^93^ into a strain homozygous for *w*^1118^, with DNA microinjection carried out at Rainbow Transgenic Flies, Inc.

### Western Blot analysis for the detection of secBirA

Ovaries from 3-day post-eclosion female flies that had been fed on yeast were dissected in PBS, moved into an Eppendorf tube and frozen in liquid nitrogen. 0-4 hour-old embryos were collected on apple juice agar plates, dechorionated, washed and frozen in liquid nitrogen. All samples were homogenized in Urea Lysis Buffer^87^ with cOmplete, EDTA-free protease inhibitor cocktail (Roche). For germline expression, ovaries and embryos were derived from females bearing *pUASp-secBirA* and *Nanos-Gal4::VP16*^58^; for follicle cells expression ovaries were obtained from females carrying the follicle cell driver *CY2-Gal4*^59^ together with *pUAST-secBirA*. Negative control extracts were generated from ovaries and embryos from females carrying only the respective Gal4 drivers. For SDS-PAGE, 150 μg of protein was loaded in each gel lane.

Following transfer to nitrocellulose blotting membrane (Amersham, Protran 0.45 μm), blots were washed briefly in Wash Buffer (25mM Tris, pH7.5, 125mM NaCl, 0.05% Tween-20) and then incubated overnight at 4°C or for 1-2 hours at room temperature in Blocking Buffer, which consisted of 25mM Tris, pH 7.5, 125 mM NaCl, 0.05% Tween-20, 5% non-fat milk and 1% BSA filtered through a paper filter (qualitative, 415, VWR). Blots were rinsed 1-2 times with Wash Buffer, then incubated overnight at 4°C with rabbit anti-BirA antibody (1 μg/ml final concentration) (Creative Diagnostics, cat # DPAB-PT1113RH) in Antibody Incubation Buffer (25mM Tris, pH 7.5, 125 mM NaCl, 0.05% Tween-20 plus 1% non-fat milk, passed through a paper filter. The blot was then rinsed 3 times in Wash Buffer followed by six 5-minute long washes, again in Wash Buffer. The blot was then incubated for 1 hour at room temperature in a solution of Peroxidase-conjugated Goat anti-Rabbit IgG (Jackson Labs) (1:5000) in Antibody Incubation Buffer. The blot was then rinsed and washed as before, and the signal was detected using the SuperSignal West Pico Kit (Thermo Scientific) and imaged using a C-DiGit blot scanner and Image Studios Software (LI-COR Biosciences).

### BirA Immunostaining

Embryos and ovaries from females expressing secBirA and PDI-GFP were collected, fixed, and immunostained using a protocol described by Coppey et al., 2008^94^ with the modification that embryos and ovaries were fixed in freshly made 4% paraformaldehyde. Rabbit anti-BirA antibody (Creative Diagnostics) was pre-absorbed against fixed embryos and used as a concentration of 1.6 μg/ml. The secondary antibody was goat anti-rabbit IgG Alexa Fluor 594 conjugate (Thermo Fisher Scientific) that was pre-absorbed against fixed embryos and used at a concentration of 2 μg/ml. The samples were mounted in Vectashield (Vector Labs) on a slide, and imaged on a Zeiss LSM 710 laser scanning confocal microscope.

### BirA activity assay

Dissected ovaries and dechorionated embryos (0-4 hour old) were homogenized in reaction buffer [(50mM Tris, pH 8.1, 500mM potassium glutamate, 0.1% Tween-20, 1X protease inhibitor cocktail cOmplete, EDTA-free (Roche)], then centrifuged at 13,500 rpm for 15 minutes at 4°C. The supernatant obtained following centrifugation was used in the assay reactions. Activity assays were carried out at 37°C for 2 hours in 36 μl of reaction buffer containing 300 μg extracted ovarian/embryonic protein, 0.325 μg/μl Maltose Binding Protein (MBP)-AviTag substrate (Avidity, L.L.C.), 8.3 mM ATP and 42 μM biotin. Negative controls consisted of extracts lacking secBirA expression and/or added MBP-AviTag. One half of the reaction was loaded into each lane. The biotinylated proteins were detected using streptavidin-HRP (Thermo Scientific) according to Hung et al., 2016^95^ with the modification of additional rinsing steps and imaged as described above.

### Visualization of biotinylated proteins in ovarian and embryonic extracts

Ovarian and embryonic extracts were generated, and SDS-PAGE gels run and blotted as described above for the BirA Western blot. Biotinylated proteins were detected using streptavidin-HRP and imaged as described above.

### Purification of biotinylated proteins

Tsl-BAP protein was isolated using the protocol described by Mayor and Peng, 2012^96^ with the modifications that all steps were carried out at room temperature, embryos were homogenized in Urea Lysis Buffer^87^, and binding to the streptavidin resin was carried out in Binding Buffer [8M urea, 200mM NaCl, 2% SDS (wt/vol), 50mM Na_2_HPO_4_, 50mM Tris, pH 8.0, protease inhibitors-cOmplete, EDTA-free (Roche)](modified from Buffer 2, from Maine et al., 2010^87^). Embryo homogenates were spun for 15-20 min at 13,200 rpm and the resulting supernatant was the starting material (SM) for the isolation. The SM was passed over a G-25 Sepharose column to remove free biotin and eluted with Binding Buffer. This was the post-equilibration (PE) sample. It was added to Pierce Streptavidin magnetic beads (Thermo Fisher Scientific) that had been pre-washed with Binding Buffer and incubated overnight in a rotater. Following this binding step, the magnetic resin was collected on the side of the tube and the buffer, the post-binding (PB) sample, was removed. Washes were carried out according to Mayor and Peng, 2012^96^. After the last wash, the resin was resuspended in 50 μl 2X Laemmli sample buffer with 8M urea (4% SDS, 20% glycerol, 200mM DTT, 125 mM Tris, pH 8.0, 8M urea). The sample was heated to 100°C for 5 minutes, spun for 5 minutes and the supernatant moved to a new tube. This was the eluate (E). Aliquots of the SM, PE, PB and E samples were subjected to SDS-PAGE and Western blotting followed by biotin detection using the Vectastain ABC AmP Reagent and Duolux chemiluminescent substrate (Vector Labs) according to the manufacturer’s directions. The blot was imaged as described above.

## Acknowledgements

We are very grateful to Drs. Paul Macdonald, Pernille Rørth, Trudi Schüpbach, and the *Drosophila* Genomics Research Center, supported by NIH grant 2P40OD010949, for providing DNA clones, *Drosophila* stocks, and *E. coli* clones. We also thank Dr. Smita Amarnath, Dr. Katie Sieverman and Ms. Emily Heines for providing experimental assistance during this study. We also acknowledge the support of Julie Hayes and the Institute for Cellular and Molecular Biology Microscopy and Imaging Facility at the University of Texas at Austin during this study.

